# The evolution of gestation length in eutherian mammals

**DOI:** 10.1101/2023.10.22.563491

**Authors:** Thodoris Danis, Antonis Rokas

## Abstract

Gestation length, or the duration of pregnancy, is a critical component of mammalian reproductive biology^1^. Eutherian mammals exhibit striking variation in their gestation lengths^2–5^, which has traditionally been linked to and allometrically scales with variation in other life history traits, including body mass and lifespan^5–8^. How the phenotypic landscape of gestation length variation, including its associations with body mass and lifespan variation, changed over mammalian evolution remains unknown. Phylogeny-informed analyses of 845 representative extant eutherian mammals showed that gestation length variation substantially differed in both whether and how strongly it was associated with body mass and lifespan across mammalian clades. For example, gestation length variation in Chiroptera and Cetacea was not associated with lifespan or body mass but was strongly associated only with body mass in Carnivora. We also identified 52 adaptive shifts in gestation length variation across the mammal phylogeny and 14 adaptive shifts when considering all three life history traits; the placements of six adaptive shifts are common in the two analyses. Notably, two of these shifts occurred at the roots of Cetacea and Pinnipedia, respectively, coinciding with the transition of these clades to the marine environment. The varying dynamics of the phenotypic landscape of gestation length, coupled with the varying patterns of associations between gestation length and two other major life history traits, raise the hypothesis that evolutionary constraints on gestation length have varied substantially across mammalian phylogeny. This variation in constraints implies that the genetic architecture of gestation length differs between mammal clades.

## Results and Discussion

### Differential associations of gestation length with body mass and lifespan across mammals

To understand the evolution of gestation length across mammals and how it has been influenced by other major life history traits, we collected data on gestation length, body mass, and lifespan for 845 eutherian mammals from the PanTheria^9^, AnAge^10^, EltonTraits^11^, and MOM^12^ databases. We then mapped the quantitative variation of all three traits onto a time-calibrated mammal phylogeny^13^. We found major differences in the patterns of variation of all three traits (**Fig. 1**). For example, gestation length and lifespan are relatively uniform within Rodentia and Perissodactyla. Within Carnivora, only Pinnipedia display elongated gestation lengths and extended lifespans, whereas taxa across the rest of Carnivora display relatively even distributions of these two traits (**Fig. 1**). Longer gestation lengths, larger body masses, and extended lifespans are also observed in Cetacea.

**Fig 1.**
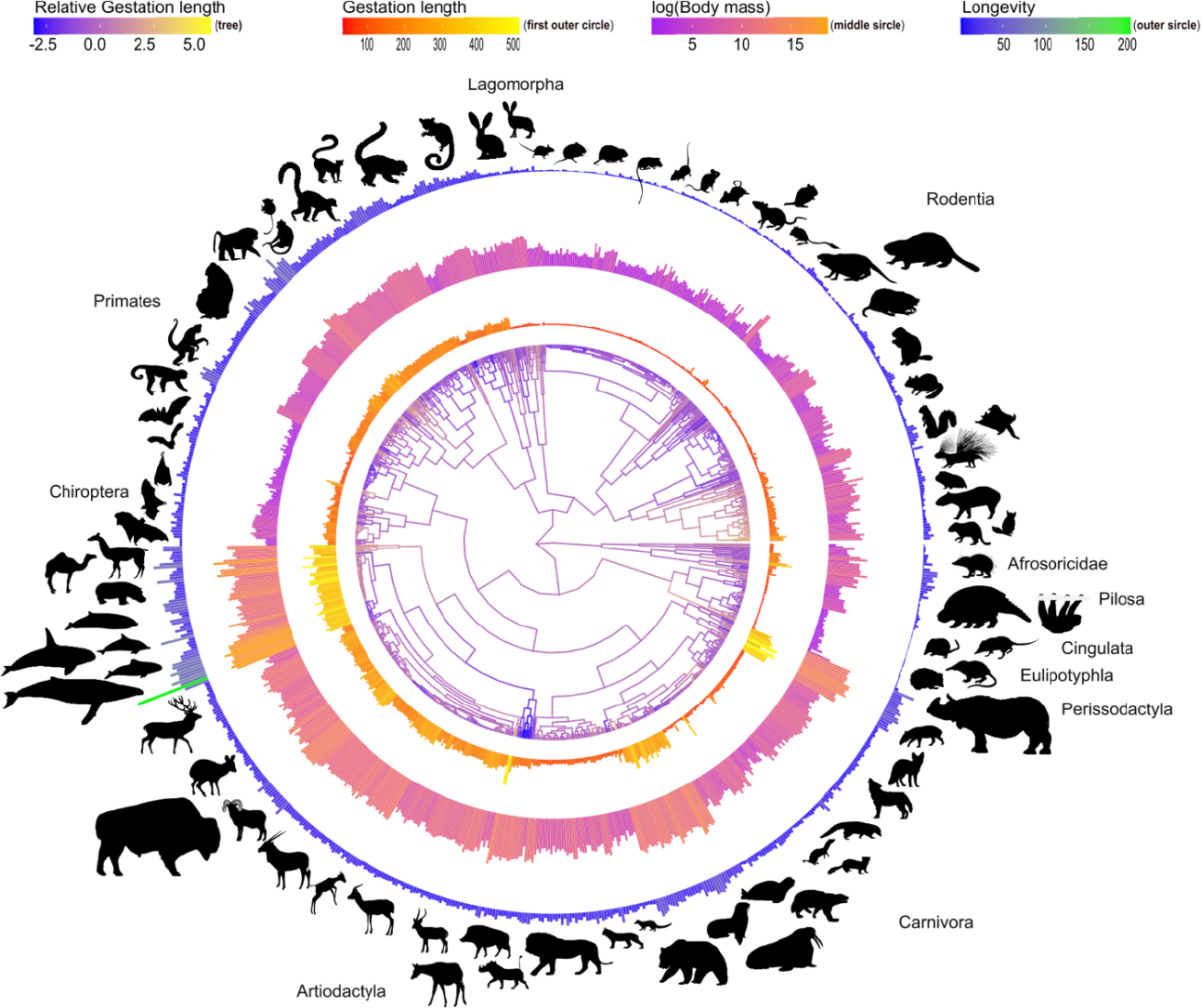
Variation in gestation length, body mass, and lifespan across 845 eutherian mammals. The graphic illustrates the relative gestation length and the absolute values of gestation length, body mass, and lifespan across the phylogeny of eutherian mammals. The branches of the phylogenetic tree illustrate relative gestation length, which was inferred by employing ancestral reconstruction on the residuals from the phylogenetic generalized least squares (PGLS) regression of gestation length ∼ body mass + lifespan + body mass * lifespan; The three outer circles depict gestation length (absolute values), body mass (log-transformed), and lifespan (absolute values), arranged from the innermost to the outermost circle; decisions on showing absolute vs. log-transformed values were for visualization purposes. Silhouette illustrations are from phylopic.org. Phylogeny is from Upham et al.^13^.

To better understand the relationship between gestation length and the other two life history traits, we performed phylogenetic regression of gestation length with body mass and lifespan as covariates using Pagel’s^14^ model. We observed significant variations in gestation length, both in terms of its presence and the strength of its association with body mass and lifespan across different mammalian lineages (**Table 1**). Specifically, there were no significant associations between gestation length and the two covariates, body mass and lifespan, or their interaction in several taxa, including Chiroptera, Artiodactyla (excluding Cetacea), Afrosoricidae, Cetacea, Pinnipedia, Lagomorpha, Pilosa, and Carnivora (**Table 1 & Fig. 2; Fig. S1**). In contrast, both New-World and Old-World monkeys (Primates) displayed significant associations between gestation length and body mass, as well as an interaction effect between the two covariates (**Table 1 & Fig. S1)**. In Perissodactyla, both covariates as well as their interaction exhibited significant interactions with gestation length. Lastly, there was a significant association between gestation length and body mass in Rodentia (**Table 1 & Fig. S1)**.

**Table 1.**
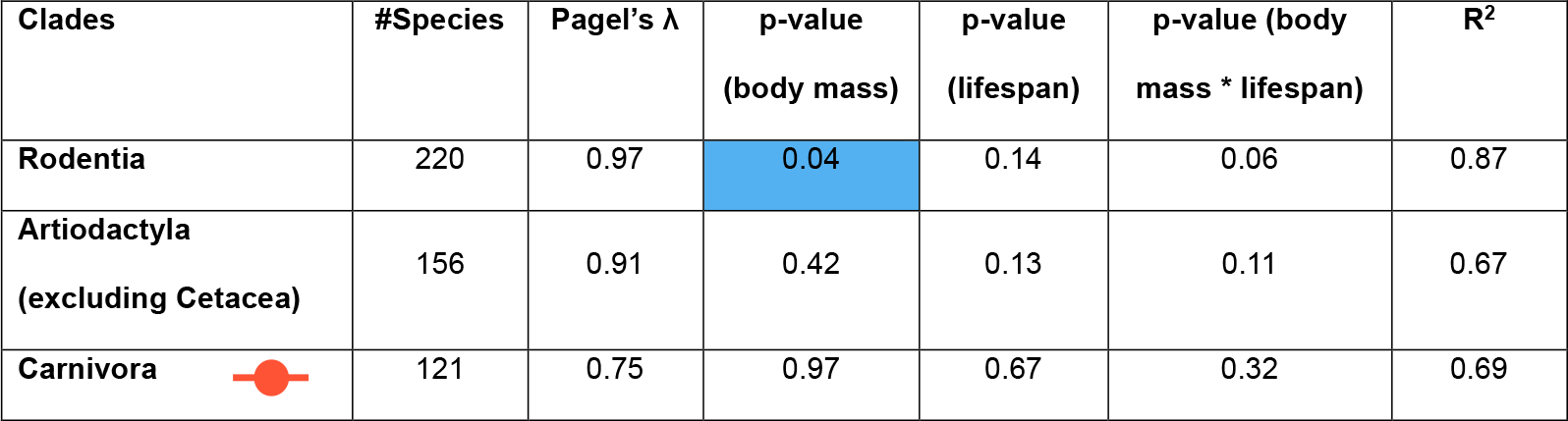
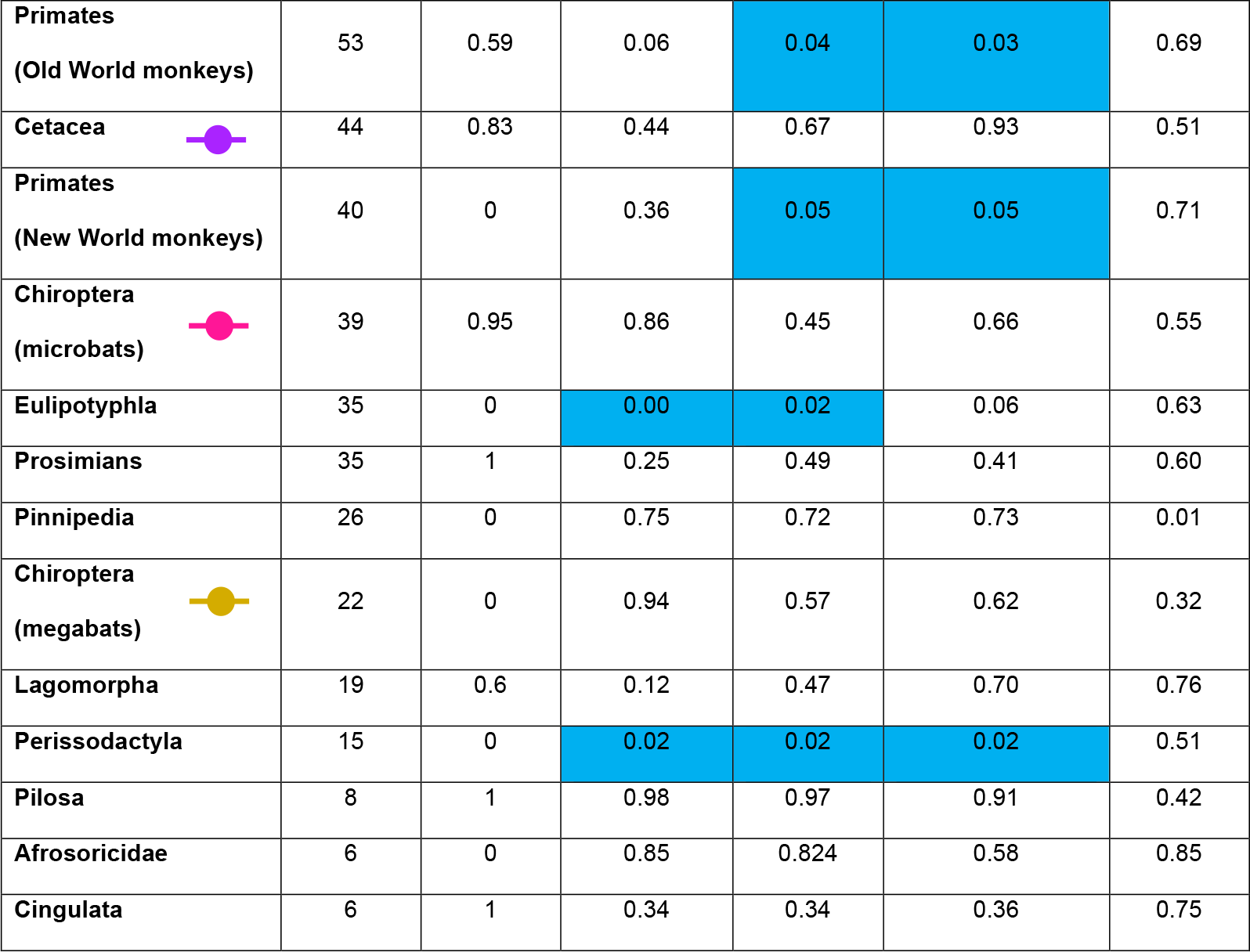
Results of Pagel’s model fitting of gestation length ∼ body mass + lifespan + body mass * lifespan. Shaded cells indicate significant p-values (< 0.05). Colored circles indicate taxa highlighted in **Fig. 2**.

| Clades | #Species | Pagel's $\lambda$ | p-value<br>(body mass) | p-value<br>(lifespan) | p-value (body<br>mass * lifespan) | R<sup>2</sup> |
| --- | --- | --- | --- | --- | --- | --- |
| Rodentia | 220 | 0.97 | 0.04 | 0.14 | 0.06 | 0.87 |
| Artiodactyla<br>(excluding Cetacea) | 156 | 0.91 | 0.42 | 0.13 | 0.11 | 0.67 |
| Carnivora | 121 | 0.75 | 0.97 | 0.67 | 0.32 | 0.69 |
| **Primates**<br>(Old World monkeys) | 53 | 0.59 | 0.06 | 0.04 | 0.03 | 0.69 |
| **Cetacea** | 44 | 0.83 | 0.44 | 0.67 | 0.93 | 0.51 |
| **Primates**<br>(New World monkeys) | 40 | 0 | 0.36 | 0.05 | 0.05 | 0.71 |
| **Chiroptera**<br>(microbats) | 39 | 0.95 | 0.86 | 0.45 | 0.66 | 0.55 |
| **Eulipotyphla** | 35 | 0 | 0.00 | 0.02 | 0.06 | 0.63 |
| **Prosimians** | 35 | 1 | 0.25 | 0.49 | 0.41 | 0.60 |
| **Pinnipedia** | 26 | 0 | 0.75 | 0.72 | 0.73 | 0.01 |
| **Chiroptera**<br>(megabats) | 22 | 0 | 0.94 | 0.57 | 0.62 | 0.32 |
| **Lagomorpha** | 19 | 0.6 | 0.12 | 0.47 | 0.70 | 0.76 |
| **Perissodactyla** | 15 | 0 | 0.02 | 0.02 | 0.02 | 0.51 |
| **Pilosa** | 8 | 1 | 0.98 | 0.97 | 0.91 | 0.42 |
| **Afrosoricidae** | 6 | 0 | 0.85 | 0.824 | 0.58 | 0.85 |
| **Cingulata** | 6 | 1 | 0.34 | 0.34 | 0.36 | 0.75 |

**Fig 2.**
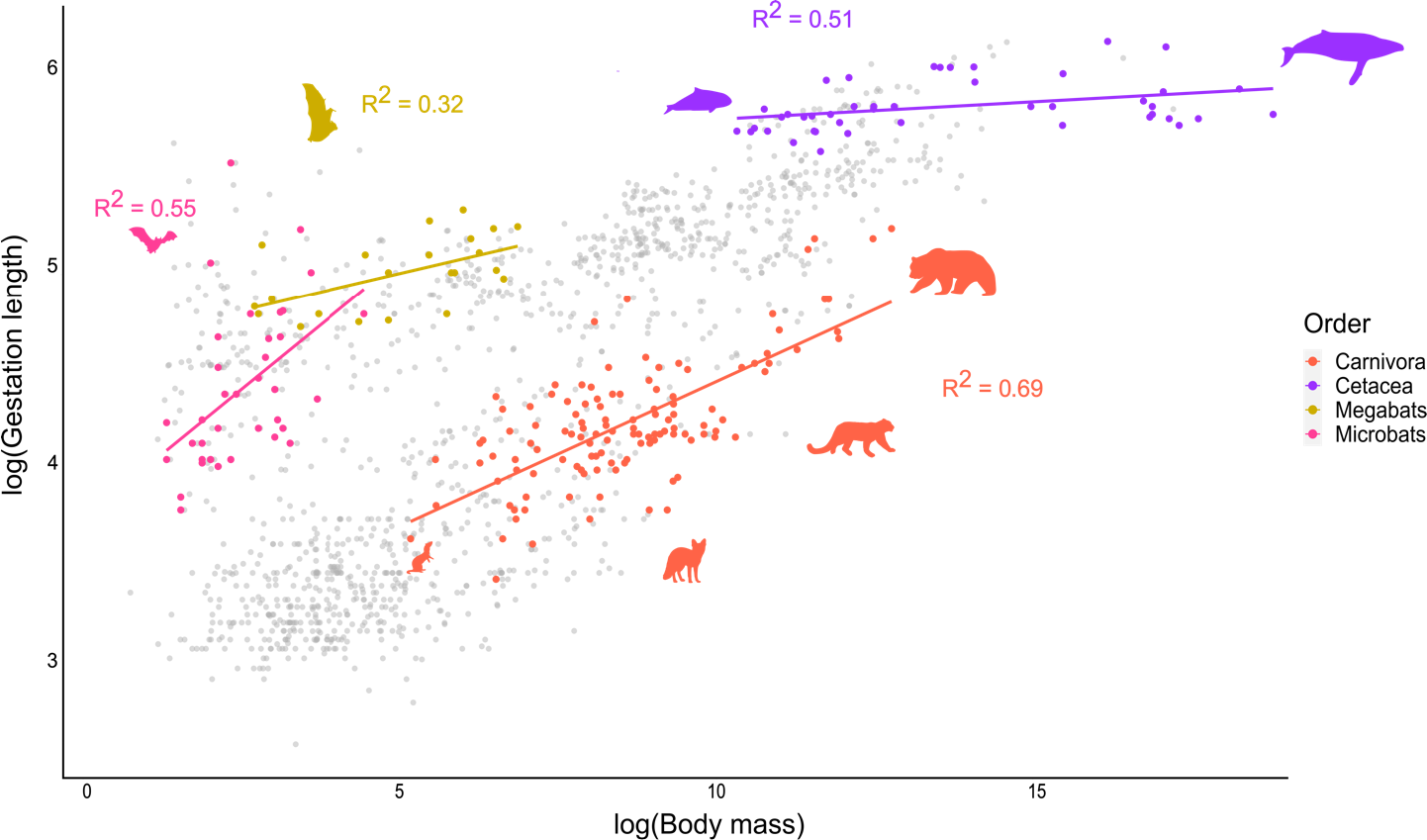
Variation in gestation length differs substantially in both whether and how strongly it is associated with body mass and lifespan across mammals. The scatterplot shows the relationship between gestation length and body mass across eutherian mammals with each dot corresponding to one of the mammalian species used in our study. Phylogenetic regression analysis using Pagel’s^14^ model was performed on three mammalian orders: Carnivora, Cetacea, and Chiroptera (megabats and microbats). Each colored data point represents a species within these orders; data points in grey correspond to species from the rest of eutherian mammals. The R^2^ values show the proportion of variance explained by the model (gestation length ∼ body mass + lifespan + body mass * lifespan) in each clade. Silhouette illustrations are from phylopic.org. The results of all eutherian taxa tested are shown in **Table 1**.

These findings challenge the notion that gestation length allometrically scales with body mass and other life history traits *across* mammals^8^, instead suggesting that different taxa have experienced differing and varying levels of selection pressure on gestation length and its linkage to other life history traits. The varying association of gestation length with other life history traits may be linked to ecological changes or historical events that differentially affected different mammalian taxa.

### Multiple evolutionary shifts in gestation length in mammalian evolution

Changes in the environment or in selective pressures can drive modifications of phenotypic traits, leading to evolutionary shifts^15^; these shifts are best interpreted as changes in trait optimal values^16^. To deepen our understanding of the evolutionary trajectory of gestation length and identify its evolutionary shifts, we first performed a univariate Bayesian analysis of gestation length. We found 52 evolutionary shifts with posterior probabilities greater than 0.25 in the evolution of gestation length across eutherian mammals. Of these 52 shifts, 29 were positive and increased gestation length, and the remaining 23 were negative and decreased gestation length (**Fig. 3**). Most of the shifts occur within mammalian orders, except for a negative shift at the root of Rodentia and Lagomorpha. The number and direction of these shifts differs across orders. For example, there are several negative shifts in the evolution of gestation length within Chiroptera, while Primates exhibit four positive shifts early in their evolutionary history. Rodentia also contain several positive and a few negative evolutionary shifts, whereas Artiodactyla have a relatively stable gestation length. Interestingly, there are positive shifts toward longer gestation lengths at the base of Pinnipedia and at the base of Cetacea, two taxa that independently transitioned to the marine environment, suggesting that some of the observed evolutionary shifts in gestation length across the mammalian phylogeny were likely closely associated with significant ecological events.

**Fig 3.**
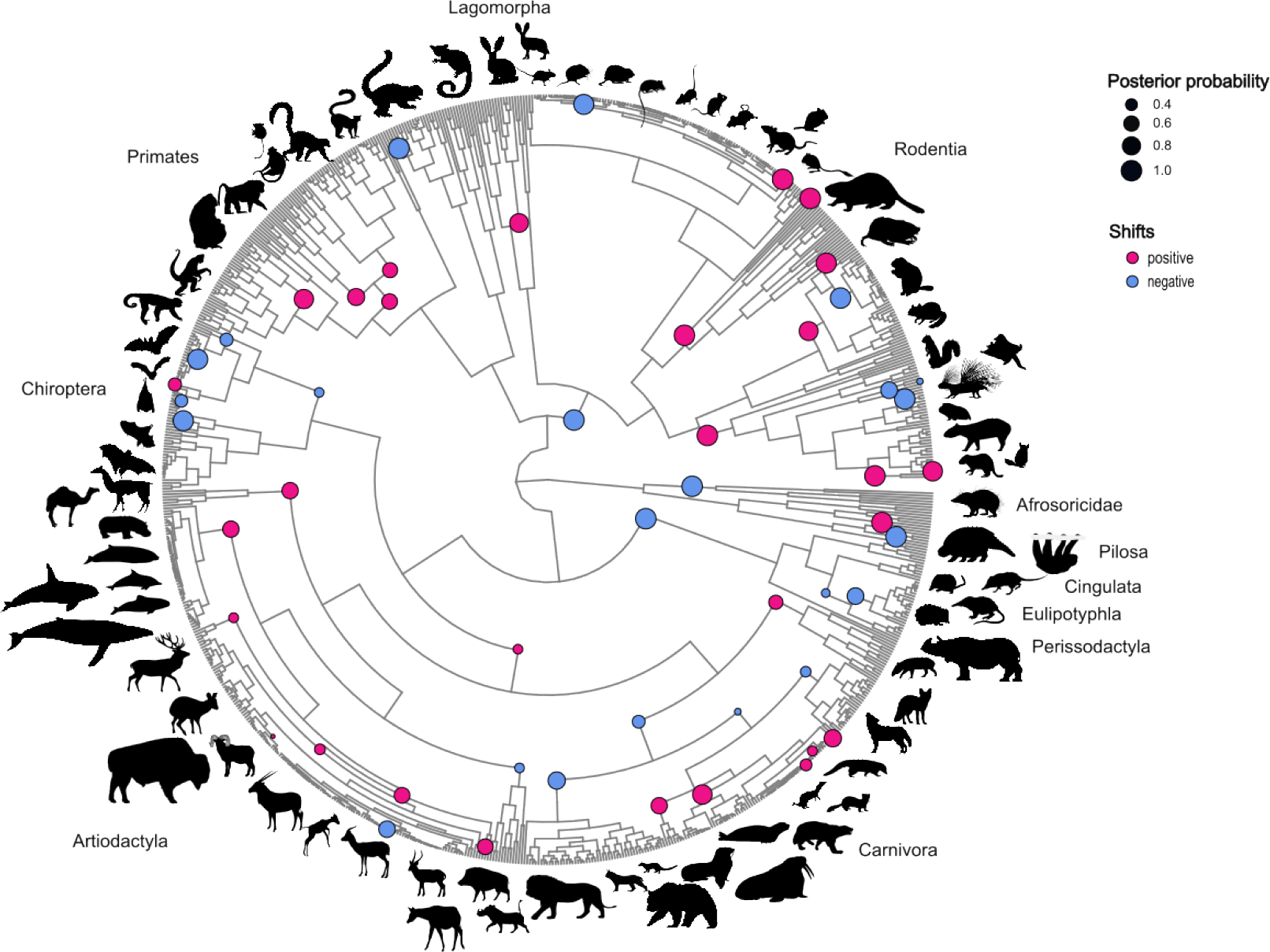
Changes in the phenotypic optimum of gestation length (evolutionary shifts) have been frequent in mammalian evolution. Circles denote the placements and magnitudes of the 52 evolutionary shifts in gestation length inferred by the Bayou Bayesian method^16^. The color of each circle corresponds to the direction of the evolutionary shift (positive shifts, n=29, dark pink; negative shifts, n=23, blue) and the size of each circle to the posterior probability of the shift; only shifts with a posterior probability greater than 0.25 are included. Silhouette illustrations are from phylopic.org. Phylogeny is from Upham et al.^13^.

Given that the evolution of gestation length is correlated with the evolution of body mass and / or lifespan in several taxa (**Table 1** and **Fig. 2; Fig. S1**), we also conducted a multivariate analysis that jointly considered the evolution of these three variables across eutherian mammals. This analysis revealed 14 shifts (**Fig. 4**), most of which occurred at or near the roots of different orders and sustained their positive trend (**Table S2**). Three of the shifts occurred at the roots of the orders Artiodactyla, Chiroptera, and Primates. Six of the 14 shifts overlap with shifts also found in our univariate analysis of gestation length (**Fig. 3**). Interestingly, two of these shared shifts occurred at the roots of the Cetacea and Pinnipedia clades and, in both cases, involved increases in all three traits, indicating a sustained evolutionary trend towards larger body masses, longer lifespans, and extended gestation periods. For Pinnipedia, body mass increased roughly 7 times as much as lifespan and ∼2.7 times as much as gestation length (estimates based on the unit changes observed in the traits’ theta values). For Cetacea, body mass increased roughly four times as much as lifespan and gestation length. There was an additional shift in our multivariate analysis within the Mysticeti, where body mass increased more than four times as much as lifespan, but gestation length remained unchanged.

**Fig 4.**
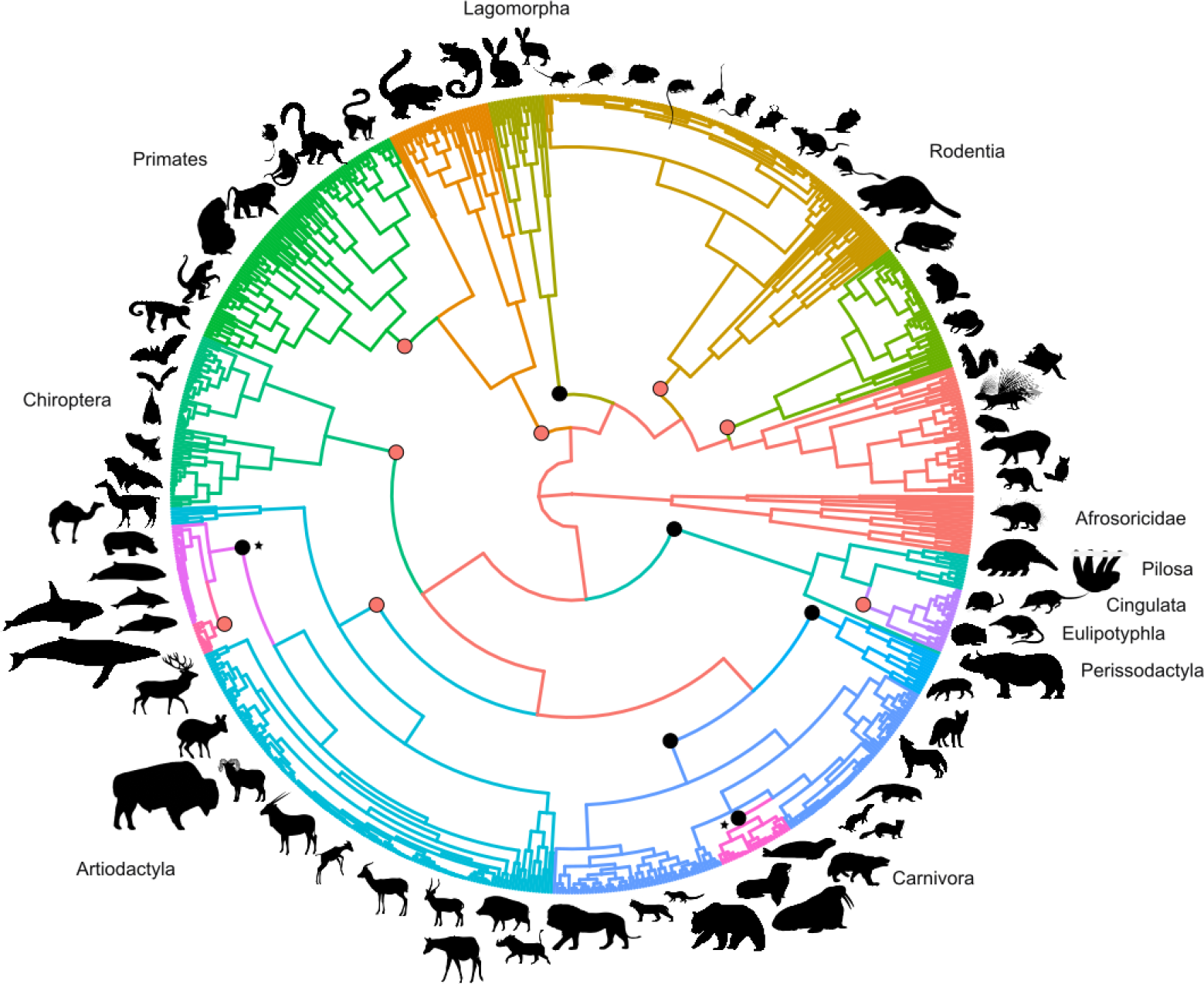
Joint consideration of gestation length, body mass, and lifespan reveals multiple evolutionary shifts during mammalian evolution. Circles denote the placements of the 14 shifts when jointly considering the evolution of gestation length, lifespan, and body mass across mammals using the PhylogeneticEM method^15^. The six evolutionary shifts common to this analysis and the analysis reported in Fig. 3 are colored black. The two asterisks correspond to the evolutionary shifts at the bases of the Cetacea and Pinnipedia clades, respectively. Branches are colored for easier visualization of the lineages that experienced evolutionary shifts. Silhouette illustrations are from phylopic.org. Phylogeny is from Upham et al.^13^.

We did not observe any evolutionary shifts in our multivariate analysis where changes in gestation length were greater in magnitude than changes in the other two traits. In general, the greatest shifts in trait values occurred for body mass, followed by gestation length and lifespan. These results suggest that the evolution of gestation length in eutherian mammals is highly interconnected with body mass and lifespan in some taxa but appears to evolve independently of the other two covariates in others^2,8,20^.

## Conclusions

By comprehensively examining the relationships between gestation length, body mass, and lifespan across 845 eutherian mammals (∼14% of extant species^21^), we reconstructed the tempo and mode of gestation length evolution. We found evidence for numerous evolutionary shifts in the gestation length optima at the origins of diverse taxa; some of these shifts were linked to changes in body mass and lifespan phenotypic optima and associated with major ecological transitions (e.g., the terrestrial to marine transition at the base of the Cetacea and Pinnipedia clades).

While our analyses yield valuable insights into the evolutionary trajectory of gestation length, we also acknowledge potential caveats and limitations. Aiming to maximize the number of mammalian species included in our study, our analyses of gestation length considered data for only two other life history traits, body mass and lifespan. There are potentially several additional life history traits that may have co-varied with gestation length, influencing its evolution, such as litter size, neonate developmental status, and mating system^1,2,22,23^. Unfortunately, data for many life history traits are scarce, so their inclusion would have dramatically reduced the number of taxa in our analyses. Additionally, we note that gestation length is not easily detectable from the fossil record, making it difficult to validate predicted evolutionary shifts by its examination.

The evolution of gestation length is associated with the evolution of body size and / or lifespan in some taxa and not associated in others, a finding that has implications for the genetic architecture of gestation length in different mammalian clades. For example, variation in gestation length is strongly associated with variation in body size in Primates. This finding is consistent with human genome-wide association studies (GWAS) of gestation length where some of the genetic variants that influence gestation length are also known to influence birth weight^24,25^. However, the lack of association between gestation length and body size in many other eutherian taxa raises the hypothesis that the genetic architecture for gestation length in these taxa will also differ. Consequently, we might expect correlational selection (i.e., selection favoring correlations between interacting traits) in taxa where gestation length coevolves with body mass and lifespan, but lack of correlational selection in lineages where the traits evolve independently of each other.

## Methods

### Data collection

We retrieved data on gestation length, lifespan, and body mass for 845 representative extant species of eutherian mammals. This information was sourced from four databases: PanTHERIA^9^, AnAge^10^, EltonTraits^11^, and MOM-Mammals^12^. To ensure the quality of our data, we only included values for adult individuals of each species and conducted a meticulous manual inspection to resolve any discrepancies. Before analysis, we natural log transformed all continuous predictor variables to reduce skewness and improve the accuracy of our findings. The study employed a consensus phylogeny obtained from the supertree reconstructed by Upham et al.^13^. This robust framework allowed us to account for evolutionary relationships among the species in our analysis. Our dataset is shown in Table S1.

### Ancestral reconstruction of relative gestation length

We extracted the residuals of gestation length while controlling for body mass and lifespan (employing the model “gestation length ∼ body mass + lifespan + body mass * lifespan”), using the function fastAnc in the package phytools v0.7^26^. We then mapped the residual values onto the mammalian phylogeny using the R packages ggtree 2.4.0^27^ and phytools 0.7^26^.

### Phylogenetic Regression

Comparisons among species violate the assumption that data points are independently drawn by a common distribution due to their shared ancestry^28^. To account for this lack of independence, we employed phylogenetic generalized least squares regression analyses (PGLS) using the ‘gls’ function in the R package nlme^29^. We used PGLS to examine the relationships of gestation length with body mass and lifespan (gestation length ∼ body mass + lifespan + body mass * lifespan) in eutherian mammals. First, we categorized the 845 eutherian mammals in our dataset into 16 clades based on their taxonomy. The 16 taxa were Afrosoricidae, Artiodactyla (excluded Cetacea), Carnivora, Cetacea, Cingulata, Eulipotyphla, Lagomorpha, Chiroptera (megabats and microbats), Primates (New World monkeys, Old World monkeys and Prosimians), Perissodactyla, Pilosa, Pinnipedia and Rodentia. Next, we fitted multiple models, and we calculated the Bayesian information criterion (BIC) for Pagel’s λ^14^ and Ornstein-Uhlenbeck (OU) model^30^. To ensure robustness, we performed 500 iterations, initializing the starting values for each model in the range of 0 to 1. Pagel’s λ^14^ model was applied using the ‘corPagel()’ function, and the Ornstein-Uhlenbeck (OU) model was applied using the ‘corMartins()’ function, both from the nlme^29^ package in R. Subsequently, we selected the best-fitting model for each mammalian group.

### Identifying evolutionary shifts in gestation length

To detect evolutionary shifts in gestation length across eutherian mammals, we used the R package Bayou v.2.2.0^16^. This tool employs a Bayesian reverse-jump MCMC approach allowing multiple optima under the OU model, identifying the number the magnitude and the location of the shifts. We implemented this approach by combining 3 parallel chains of 5 million iterations, with a burn-in proportion of 0.3. We allowed only one shift per branch and the total number of shifts was calculated based on the conditional Poisson prior with a mean equal to 2.5% of the total number of branches in the tree and a maximum number of shifts equal to 5%, following the authors’ recommendations (Uyeda et al., 2017). For all the other parameters, we used the recommended distributions in the publicly available tutorial (https://github.com/uyedaj/bayou/blob/master/tutorial.md). The MCMC was initialized randomly selected parameters for the first 1000 generations to improve the fit of the model. Finally, we ensured that independent chains had converged on similar regions in the parameter space by Gelman’s^33^ R for log likelihood,σ^2^, and α (**Fig. S2**).

### Evolutionary shifts of gestation length, body mass and lifespan across Eutherian mammals

To investigate evolutionary shifts in the evolution of all three life history traits across the eutherian mammal phylogeny, we used the PhylogeneticEM^15^ R package. This method infers evolutionary shifts in multivariate correlated traits on phylogenies via an OU process. Shift positions were estimated using the Expectation-Maximization (EM) algorithm, considering varying numbers of unknown shifts, and the optimal number of shifts was determined using a lasso-regression model selection procedure. All parameters were kept at their default settings except for the maximum number of shifts, which was set to 18.

## Acknowledgements

We thank Luis Muglia, Kristina Kverková, Graham Slater, Paul Bastide and members of the Rokas Lab, especially Dana Lin and Kyle David, for feedback and support on this project. This work was supported by the Graduate Program in Biological Sciences at Vanderbilt University, the Burroughs Wellcome Fund, and the March of Dimes Prematurity Research Center Ohio Collaborative. Research in A.R.’s lab is also supported by the National Science Foundation (DEB-2110404) and the National Institutes of Health / National Institute of Allergy and Infectious Diseases (R01 AI153356). This work was conducted in part using the resources of the Advanced Computing Center for Research and Education at Vanderbilt University.

