## Supplementary Figures and Tables for "The evolution of gestation length in eutherian mammals"

ORCIDs: Thodoris Danis: 0000-0002-0862-4972

Antonis Rokas: 0000-0002-7248-6551

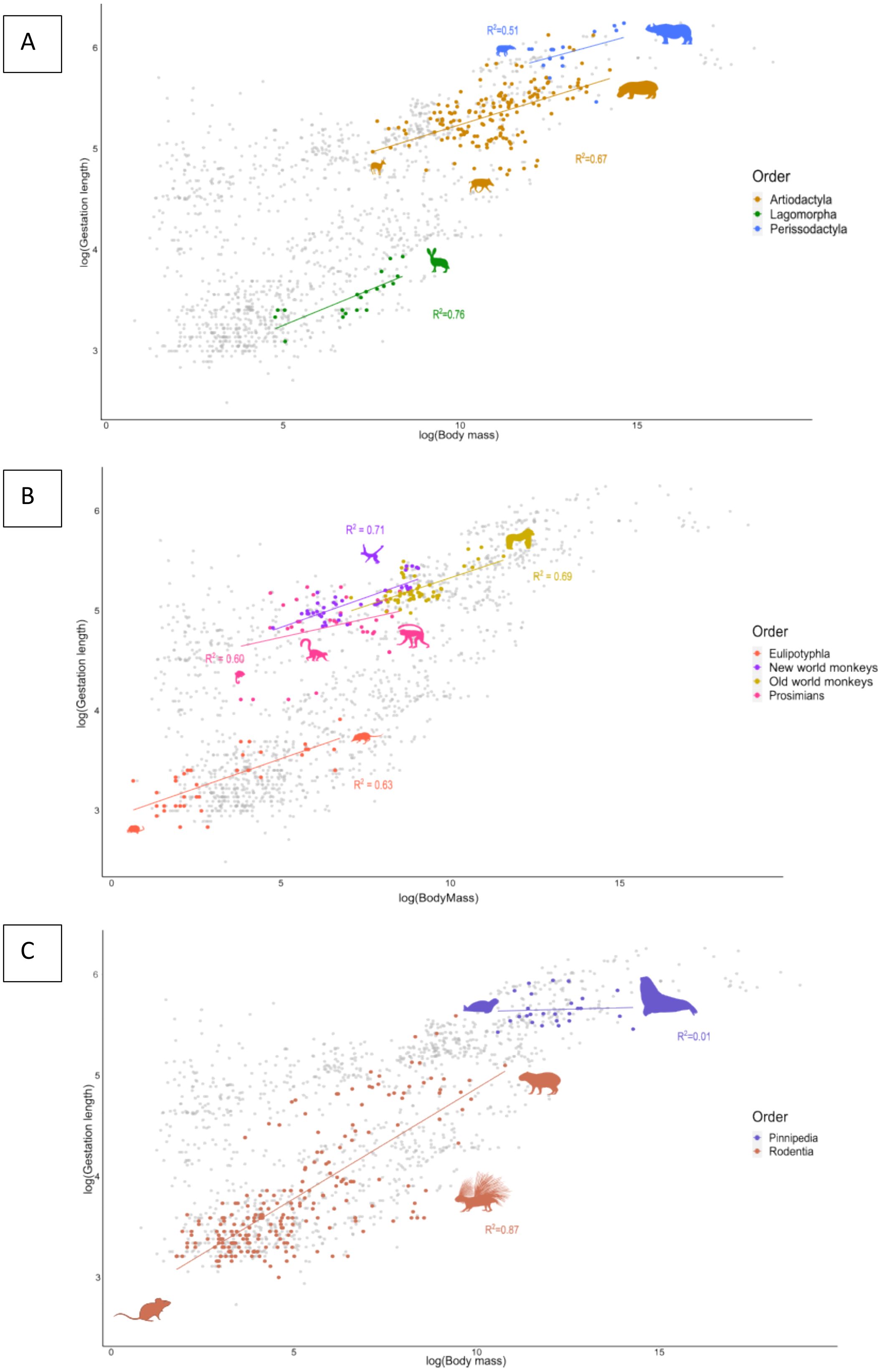
A
R²=0.51
R²=0.67
R²=0.76
log(Gestation length)
log(Body mass)
Order
Artiodactyla
Lagomorpha
Perissodactyla
B
R² = 0.71
R² = 0.69
R² = 0.60
R² = 0.63
log(Gestation length)
log(BodyMass)
Order
Eulipotyphla
New world monkeys
Old world monkeys
Prosimians
C
R²=0.01
R²=0.87
log(Gestation length)
log(Body mass)
Order
Pinnipedia
Rodentia

**Fig S1. Variation in gestation length differs substantially in both whether and how strongly it is associated with body mass and lifespan across mammals.** Scatterplots illustrating the relationship between gestation length and body mass across various eutherian mammals; each scatterplot highlights different taxonomic groups. Each dot represents a mammalian species included in our study. Phylogenetic regression analysis was performed using the Pagel's[1] model. **A.** Scatterplot highlighting the orders Artiodactyla (excluding Cetacea), Lagomorpha, and Perissodactyla. **B.** Scatterplot highlighting the orders Pinnipedia and Rodentia. **C.** Scatterplot highlighting the orders Eulipotyphla and Primates (Prosimians, New and Old-world monkeys). Colored data points represent species within these groups, while grey data points correspond to species from the rest of eutherian mammals. The $R^2$ values indicate the proportion of variance explained by the model (Gestation length ~ body mass + lifespan + body mass * lifespan) within each group. Silhouette illustrations are sourced from phylopic.org. A comprehensive table of results for all eutherian taxa examined can be found in **Table 1 of the main text**.

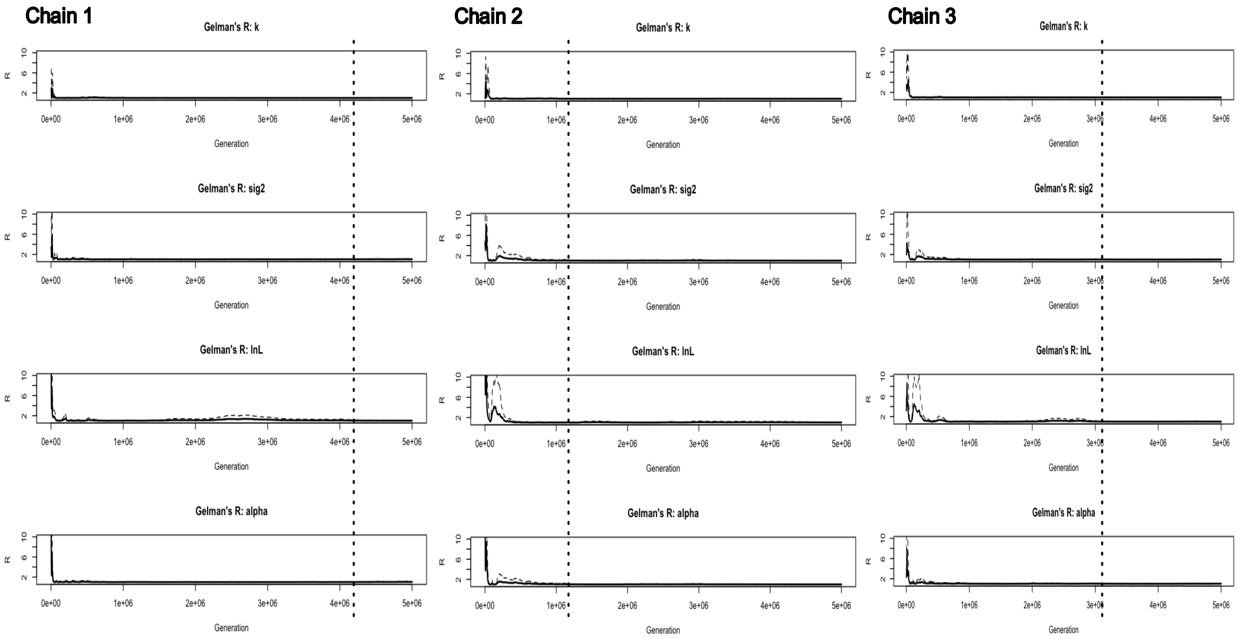

**Fig S2. Evidence of convergence in Monte Carlo chains used in the Bayesian Bayou[2] framework.** Convergence was achieved for three independent chains of $5*10^6$ iterations, as indicated by Gelman's[3] statistic for kappa, $\sigma^2$, log likelihood, and alpha ($\alpha$). The dashed line marks the point where these chains converged.

**Table S1. Results from the joint consideration of gestation length, body mass, and lifespan using the PhylogeneticEM[4] software.**

| **Shifts** | | | | |
|---|---|---|---|---|
| | **Nodes** | **Body mass** | **Gestation length** | **Longevity** |
| | **1087** | -0.684469 | 0.214965 | 0.485801 |
| | **938** | -2.973135 | -1.264069 | -0.841769 |
| | **1069** | -0.296652 | -1.099038 | -0.472871 |
| | **894** | -1.578072 | -1.042574 | -0.159839 |
| | **1122** | 1.709247 | 0.317992 | 0.413528 |
| | **1216** | -3.883559 | -0.027166 | 0.172693 |
| | **1637** | -2.010477 | -1.046395 | -0.783768 |
| | **1277** | 3.67008 | 0.734048 | 0.455924 |
| **Perissodactyla** | **1623** | 5.849071 | 1.323956 | 1.161076 |
| | **1477** | 1.577359 | -0.361292 | 0.431192 |
| | **1640** | -3.06394 | -0.439996 | -0.948205 |
| **Cetacea** | **1285** | 2.39207 | 0.543303 | 0.653471 |
| **Pinnipedia** | **1537** | 3.538727 | 1.324626 | 0.463173 |
| **Mysticeti** | **1319** | 3.891177 | -0.033671 | 0.837887 |
| **Optimal values (θ)** | | | | |
| | **Nodes** | **Body mass** | **Gestation length** | **Longevity** |
| | **1087** | 6.570128 | 4.847101 | 2.929979 |
| | **938** | 4.281462 | 3.368067 | 1.60241 |
| | **1069** | 6.957945 | 3.533097 | 1.971307 |
| | **894** | 5.676525 | 3.589562 | 2.284339 |
| | **1122** | 8.279375 | 5.165093 | 3.343507 |
| | **1216** | 3.371038 | 4.604969 | 2.616872 |
| | **1637** | 5.24412 | 3.585741 | 1.660411 |
| | **1277** | 10.92468 | 5.366184 | 2.900103 |
| **Perissodactyla** | **1623** | 13.103668 | 5.956092 | 3.605255 |
| | **1477** | 8.831955 | 4.270844 | 2.875371 |

| | | | | |
|---|---|---|---|---|
| | **1640** | 2.180181 | 3.145745 | 0.712206 |
| **Cetacea** | **1285** | 13.316747 | 5.909487 | 3.553573 |
| **Pinnipedia** | **1537** | 12.370682 | 5.59547 | 3.338544 |
| **Mysticeti** | **1319** | 17.207924 | 5.875816 | 4.391461 |

**Table S2.** Dataset of 845 eutherian mammal species and their values for the three life history traits of body mass, gestation length, and lifespan. This table will be made publicly available upon manuscript acceptance.
